# Effects of Pathological Mutations on the Prion-Like Polymerisation of MyD88

**DOI:** 10.1101/351726

**Authors:** Ailís O’Carroll, Brieuc Chauvin, James Brown, Ava Meagher, Joanne Coyle, Dominic Hunter, Akshay Bhumkhar, Thomas Ve, Bostjan Kobe, Emma Sierecki, Yann Gambin

**Affiliations:** EMBL Australia Node in Single Molecule Science, University of New South Wales, Kensington, NSW 2052, Australia.; Institute for Molecular Bioscience, University of Queensland, Brisbane, QLD 4072, Australia.; Institute for Glycomics, Griffith University, Southport, QLD 4222, Australia; School of Chemistry and Molecular Biosciences, and Australian Infectious Diseases Research Centre, University of Queensland, Brisbane, QLD 4072, Australia.

## Abstract

A novel concept has emerged whereby the higher-order self-assembly of proteins provides a simple and robust mechanism for signal amplification. This appears to be a universal signalling mechanism within the innate immune system, where the recognition of pathogens or danger-associated molecular patterns need to trigger a strong, binary response within cells. Previously, multiple structural studies have been limited to single domains, expressed and assembled at high protein concentrations. We therefore set out to develop new in vitro strategies to characterise the behaviour of full-length proteins at physiological levels. In this study we focus on the adaptor protein MyD88, which contains two domains with different self-assembly properties: a TIR domain that can polymerise similarly to the TIR domain of Mal, and a Death Domain that has been shown to oligomerise with helical symmetry in the Myddosome complex. To visualize the behaviour of full-length MyD88 without purification steps, we use single-molecule fluorescence coupled to eukaryotic cell-free protein expression. These experiments demonstrate that at low protein concentration, only full-length MyD88 forms prion-like polymers. We also demonstrate that the metastability of MyD88 polymerisation creates the perfect binary response required in innate signalling: the system is silenced at normal concentrations but upstream signalling creates a “seed” that triggers polymerisation and amplification of the response. These findings pushed us to re-interpret the role of polymerisation in MyD88-related diseases and we studied the impact of disease-associated point mutations L93P, R196C and L252P/L265P at the molecular level. We discovered that all mutations completely block the ability of MyD88 to polymerise. We also confirm that L252P, a gain-of-function mutation, allows the MyD88 mutant to form extremely stable oligomers, even when expressed at low nanomolar concentrations. Thus, our results are consistent with and greatly add to the findings on the Myddosomes digital ‘all-or-none’ responses and the behaviour of the oncogenic mutation of MyD88.

## Introduction

In the innate immune system, dedicated germline encoded receptors known as Pattern Recognition Receptors (PRRs) recognise pathogens from all major classes of invading microorganisms, as well as other endogenous danger associated molecular patterns. The Toll Like Receptors (TLRs) are a major family of PRRs whose signalling pathways culminate in the activation of transcription factors that mediate innate immune responses and therefore have a crucial regulatory function in maintaining health and eradicating disease^1^. TLRs recruit different combinations of the five main adaptor proteins (TRIF, TRAM, SARM, Mal and MyD88)^2^. MyD88 is the least polymorphic adaptor and has evolved under purifying selection, confirming its role as an essential and non-redundant protein in host survival^3^. This implies a crucial role in signalling. Patients with MyD88 mutations within MyD88 such as theL93P andR196 point mutations present a primary immunodeficiency syndrome characterized by greater susceptibility to pyogenic Gram-positive bacterial infections^4,5 6^, which often result in life-threatening bacterial infections. Interestingly, somatic mutations in MyD88 have also been found that contribute to human malignancies for both chronic lymphocytic leukaemia and more commonly, diffuse large B cell lymphoma^7^. In particular, the L252P point mutation (also referred to as L265P) was discovered to be driving the promotion of malignant cell survival in the lymphomas present in many cancer patients^4, 16^.

MyD88 is recruited to TLR4 through another adaptor protein, Mal. Upon receptor oligomerisation^2^, Mal acts as a nucleation platform for the downstream recruitment of MyD88 through homotypic interactions between their respective Toll-Interleukin Receptor (TIR) domains ^8^. MyD88 also possesses a Death Domain (DD) (1-110 a.a) that binds to and recruits the downstream kinases IRAK2 and IRAK4^9^. The death domains of MyD88, IRAK2 and IRAK4 co-assemble into the well-defined “Myddosome”. The structure of the Myddosome was solved by crystallography^9–11^ and shows a helical organization of 6-8 DD of MyD88, 4 IRAK2 and 4 IRAK4 DD molecules^9^. The spatial localization of the TIR domain of MyD88 and its role in the Myddosome assembly could not be determined in these studies as only the DD of MyD88 was used. In parallel, recent studies demonstrate the role of the TIR domain in the self-assembly of MyD88. Ve et al. discovered that the recombinant TIR domains of Mal and MyD88 alone can self-organize into helical filaments and have solved the structure of the assembly by cryo-electron microscopy^12^. These TIR filaments of both Mal and MyD88 have self-replicating propensity, joining the growing list of “prion-like” polymers found within the proteins of the immune system, namely RIPK1, RIPK3^13^, MAVS (mitochondrial antiviral-signalling protein) and ASC (Apoptosis-Associated Speck-Like Protein Containing CARD^9, 10, 12^.

From these discoveries, a novel paradigm is emerging, recently termed ‘signalling by cooperative assembly formation’ or SCAF^14^, whereby the response of the innate immune system is driven through the polymerisation of adaptor proteins, creating a highly non-linear amplification of the signal. Intriguingly, many adaptors with prion-like properties contain two domains, both with self-assembly properties. For example with ASC, the PYD and CARD domains can both form filaments separately, and although the structure of the full-length ASC protein upon polymerisation is still unknown^15, 16^. Similarly, for MyD88, both DD-Myddosome and TIR filament structures have been solved but the interplay between the two domains and their contribution to both processes is unknown.

To characterize the contribution of self-association propensities from the TIR and DD domains to full-length MyD88, we used a combination of single-molecule fluorescence microscopy techniques and ‘*in-vitro*’ protein expression. This enabled us to characterize the behaviour of full length MyD88 compared to that of the individual domains, by precisely measuring their degree of oligomerisation or fibrillation, at low concentrations and without purification. In the same system, we studied the effect of 3 disease-associated point mutations, one mutation in the DD (L93P) and two within the TIR domain (R196C and L252P). We showed that all three mutations have the ability to abrogate the polymerisation of MyD88 into filaments. Our experiments also revealed an unexpected behaviour for the L252P mutant, which dramatically enhances MyD88 self-association (at lower concentration) into small and well-defined oligomers, creating extremely stable assemblies.

## RESULTS

Our aim was to look at the self-assembly of full-length MyD88 and characterise the contribution of both protein domains in this process. Full length MyD88 is difficult to express and purify recombinantly in E. Coli, presumably due to its polymerisation propensity. Here we express the proteins *in-vitro* at controlled low concentrations and study their self-association in undisturbed samples using single-molecule counting techniques. More precisely, we use an *in-vitro* translation system derived from *Leishmania tarentolae* ^17^ (*Leishmania tarentolae* extract, LTE). This eukaryotic system enables the rapid production of proteins (typically within 2hours) and the analysis of protein-protein interactions in a system orthogonal to the human proteome. By controlling the concentration of DNA priming the system, we can tune the final expression levels of proteins and co-express proteins at controlled ratios. We have tested this combination on a variety of biological systems over the years, and demonstrated that the flexibility of cell-free protein expression is a great asset to study protein aggregation and prion-like fibrillation^15, 18–21^.

To determine the aggregation and oligomerisation propensity of proteins, we developed diverse “counting” methods based on single-molecule fluorescence techniques. In single molecule fluorescence, rare protein complexes can be easily detected in a background of monomers and their size can be evaluated by simply counting the number of fluorophores present in each complex. As we demonstrated recently in our study of the prion-like behaviour of ASC, these counting methods are well suited to study heterogeneous processes of protein oligomerisation and polymerisation^15^. To visualize MyD88 and its mutants, the proteins are expressed as fusion with genetically encoded GFP or mCherry fluorophores, and can be measured directly upon expression without further labelling, purification or enrichments steps. The levels of fluorescence obtained provide a direct readout of protein expression levels, after careful calibration with GFP / mCherry protein controls.

First, constructs containing the TIR domain alone, death domain alone (1-117 a.a to include crucial part of ID) or full-length MyD88 fused to GFP at the N-terminus were expressed in LTE. Upon expression, proteins were diluted 10 times (to concentrations ranging from 0-300nM) and placed onto a confocal microscope. A 488 nm laser was focused into the sample, creating a small focal volume, through which proteins freely diffuse due to Brownian motion. In this range of concentration multiple fluorophores are always present in the focal volume and as proteins constantly exchange in the detection volume, we ultimately interrogate a large number of proteins. Fluctuations of fluorescence intensity were recorded using high speed single photon counters and typical fluorescence time traces, shown in Figure 1A were obtained.

**Figure 1:**
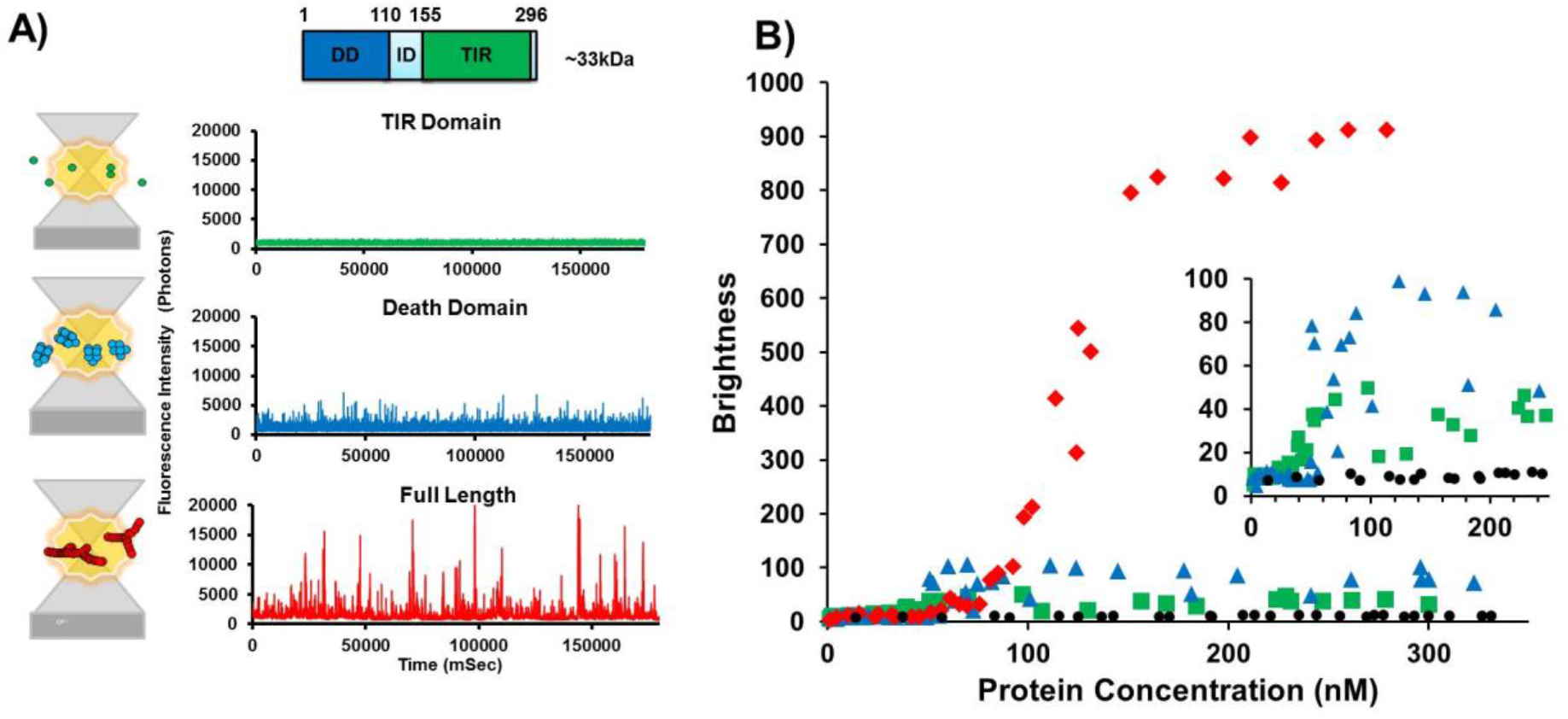
The domains of MyD88 exhibit different oligomerisation propensities, with only the full-length protein forming polymers. (A) Schematic diagram of single-molecule counting experiments, demonstrating the distinction between the oligomeric protein sizes measured, as the green fluorescently-tagged protein complexes excited by a 488 nm laser diffuse freely in and out of the focal volume. The schematic diagrams reflect the fluorescence time-traces obtained. The diffusion of an oligomer equates to the same number of fluorophores moving through the confocal volume, creating a burst of fluorescence in the time-trace being directly proportional to the size of the oligomer. For GFP-tagged MyD88 TIR domain, small fluctuations in intensity are recorded around the average fluorescence value, as expected for an extremely low order oligomer such as a dimer (for example, if 20 proteins are detected simultaneously, the exit/entry of a single protein causes a decrease/increase of signal of only 5%). The GFP-tagged MyD88 DD shows larger bursts of fluorescence correlating with these death domains forming higher-order oligomeric complexes. As seen by the fluorescent time traces, N-terminally GFP tagged full-length MyD88 shows extremely large filamentous polymers of MyD88 diffusing through the confocal volume. (B) The B parameter (Brightness) correlating with number of oligomers detected in typical time-traces as a function of protein concentration (nM), for the TIR domain (green), DD (blue) and wild-type full length MyD88 (red). Protein concentrations range from 0 to 320nM.

### At low concentrations, both the TIR and death domains are required for efficient polymerisation of MyD88

The fluorescence time-traces obtained for the full-length and separate domains of MyD88 (Figure 1A) exhibit very distinct characteristics. For TIR domain alone (Figure 1A, in green), the fluctuations of intensity around the average value are limited (± 500 photons/ms), however for the death domain (Figure 1A, in blue) small bursts of intensities (> 1500 photons/ms above background) can be detected. These fluorescence peaks correspond to entries of single protein complexes, increasing the local number of proteins for a brief period of time. The amplitude of the deviations from average is linearly linked to the number of proteins co-diffusing in a single complex, and the duration of the burst is linked to the physical size of the diffusion complex. For the full-length MyD88, we observe extremely bright and long-diffusing bursts of fluorescence, as shown in red.

The simplest analysis to quantify the presence of protein complexes is to calculate the average brightness of the diffusing species. The brightness parameter B is calculated from the intensity values measured as

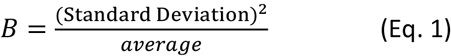

The main advantage of this B parameter is that it is independent from protein concentrations, so one can easily visualize self-association as a function of expression levels. Variation of the final levels of protein expression was achieved by using serial dilutions of the priming DNA template in our cell-free expression system, with lower DNA concentrations resulting into lower protein concentrations. For each experiment, fluorescence times were recorded and the brightness parameter was calculated and plotted as a function of protein expression concentration. As shown in Figure 1B, all three constructs display self-association compared to the GFP control. TIR alone forms small oligomers which typically contain 3 GFP fluorophores, and the concentration-dependence shows that TIR self-assembles around 50 nanomolar. The DD only construct forms larger assemblies by itself, containing approx. 8-10 GFP molecules. The concentration-dependence shows a sharp increase into self-assembly at 60 nanomolar. The full-length protein displays much higher brightness values, suggesting at least 80 GFP co-diffusing. The apparent Kd of self-assembly is displaced compared to DD and TIR alone, with 50% assembly reached at 120 nanomolar.

To evaluate the size of the oligomers and assemblies more precisely, we used Photon Counting Histograms (PCH) analysis to determine the stoichiometry of the complexes. We have described the PCH method in detail in previous publications^15, 19, 22^, and will provide a brief description here of the principle and analysis. To perform Photon Counting, the sample (expressed at the highest DNA loading) is diluted so that individual peaks can be fully resolved. The dilution of the sample to picomolar concentration ensures that the probability of observing multiple complexes at the same time is extremely low, and hence one can attribute single peaks of fluorescence to single, individual complexes. In that case, the maximal number of photons emitted by a complex is directly linked to the number of fluorophores within the complex, thus each peak can be quantified with the “single molecule ruler” of monomeric GFP. Simply put, as we know that a single monomeric GFP can emit a maximum of 110, fluorescence bursts with >10,000 photons observed for MyD88 corresponds to >100 proteins co-diffusing. Based on GFP, we estimate that the TIR domain forms a trimer while the DD is mainly forming octamers (Figure 2A). These differences are confirmed by Fluctuation Correlation Spectroscopy (FCS) that gives information on the average physical size of the diffusing particles. As shown in Figure 2B-C, the MyD88 full-length form complexes that diffuse > 100-fold slower than single GFPs. In comparison, the TIR domain assembly diffuses approximately 2-fold slower than GFP monomers, consistent with the idea that it is trimeric while the DD assemblies demonstrate approximately 8-fold slower diffusion when compared to the GFP monomer.

**Figure 2:**
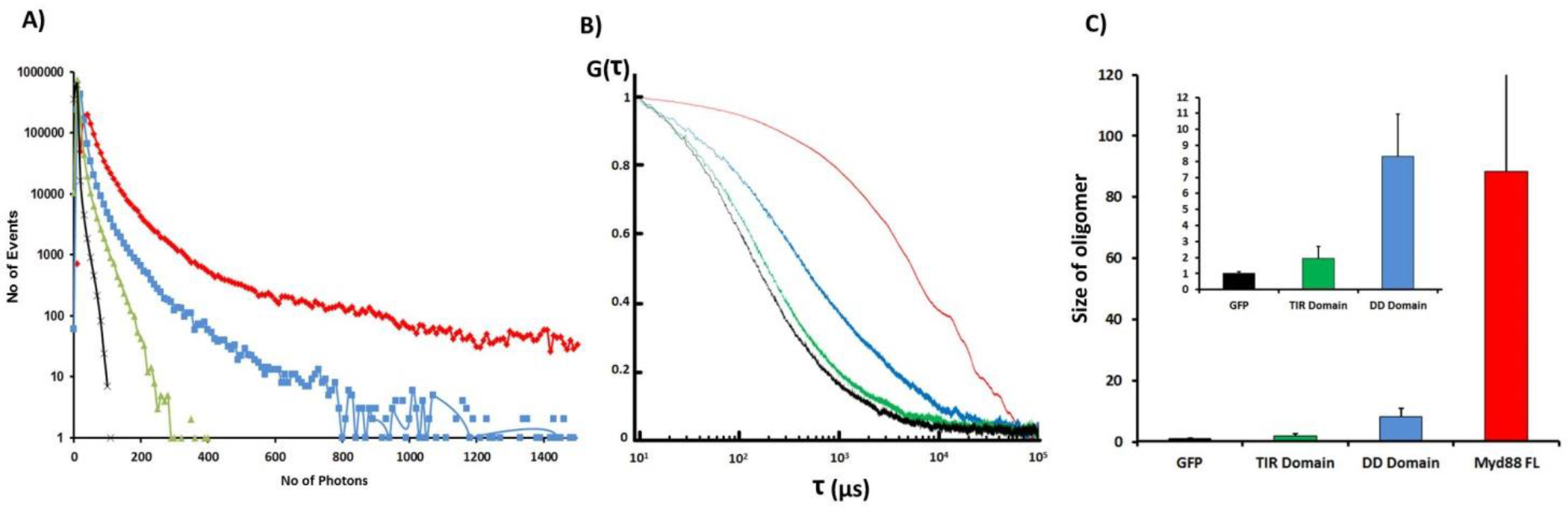
Calculation of TIR domain, DD alone and MyD88 full-length size based on single molecule techniques. (A) PCH fluorescence intensity histograms with a binning in 10 μs. The number of events at the various fluorescence intensities allows us to clearly demonstrate the divergences in size from the background GFP level. This allows us to define the approximate oligomerisation status of the full length proteins and domains. (B) FCS data in solution. Control based on GFP monomer (black). Correlation curves obtained for the TIR domain (green), DD (blue) and MyD88 full-length (red). A clear shift in diffusion time can be seen between the full-length and the domains alone. (C)Brightness histogram normalised by the GFP control (black), therefore allowing the calculation of approximate size of the oligomeric species.

Overall, this data is consistent with the idea that MyD88 full-length is more prone to fibrillation than the isolated domains that tend to form only low order oligomers. This is independent of the constructs of full-length MyD88 used (i.e. tagged with C- and N-terminal mCherry and His-tagged constructs) as all demonstrated the same behaviour independently of their tag (Data not shown). Comparisons of the brightness profiles with PCH and FCS data obtained with TIR alone and purified GFP-foldon (a control trimeric GFP-tagged protein) confirm that in our hands, the TIR domain species are trimeric (Figure 2A-C). This behaviour was consistent no matter what concentrations of the protein were expressed (Figure 1B). The absence of large events is consistent with the findings by Ve et al. that MyD88 TIR alone does not spontaneously polymerise^12^. Similarly, comparisons of the brightness profiles, PCH and FCS and purified GFP-foldon data obtained when the MyD88 DD alone is expressed, demonstrates that it too consistently forms small oligomers, more specifically octamers based on PCH analysis (Figure 2A-C). This data complements the previous Myddosome data that shows assemblies of 6-8 MyD88 DDs^23^. Our results are also consistent with the recent single molecule fluorescence microscopy studies by Latty et al. ^23^ demonstrating the formation of both smaller (6 MyD88 complexes approximately) and ‘super’ Myddosomes at the cell surface. Overall, the different domains exhibit unique oligomerisation propensities, with both domains presence necessary but not sufficient for the formation of polymers.

### MyD88 aggregation is a concentration-dependent, self-templated polymerisation event

Figure 1B showed that the aggregation of MyD88 full-length was a concentration-dependent process and revealed a sharp transition in behaviour at around 120 nM.

To verify that this self-assembly was a true dynamic process, we used a 2 colour seeding assay (Figure 3A). Briefly, full-length MyD88 tagged with mCherry was expressed at high concentrations (250nm) to allow the formation of filaments. These objects were enriched by gentle spinning and sonication before being added to solutions containing MyD88 tagged with GFP expressed across a range of concentrations, as previously described. Self-templating is determined by using two-colour single particle coincidence. This time, two lasers (488 nm and 546 nm) are focused on the same focal volume, allowing both mCherry and GFP-tagged proteins to be detected at the same time. A typical fluorescence time trace demonstrating two colour-coincidence experiments with MyD88 is shown in Figure 3A. The initial expression of GFP tagged MyD88 is at subcritical concentrations in the absence of mCherry MyD88 sonicated filaments (“seeds”): a GFP trace showing little fluctuation is obtained indicating that MyD88 is monomeric at this concentration. The mCherry-tagged MyD88 seeds are then added to the mixture and are detected in the mCherry channel. If GFP is recruited to the mCherry seeds, this results in the presence of coincident bursts of fluorescence in both channels. Indeed, within 20 seconds, large bursts of fluorescence in the GFP channel are seen and they predominantly coincide with the presence of mCherry peaks, indicating that MyD88-GFP is recruited to the MyD88 mCherry seeds (Figure 3B). Moreover, with time, the events detected in GFP become brighter than the ones in Cherry, indicating that the GFP-tagged MyD88 grow off the MyD88 Cherry seeds.

**Figure 3.**
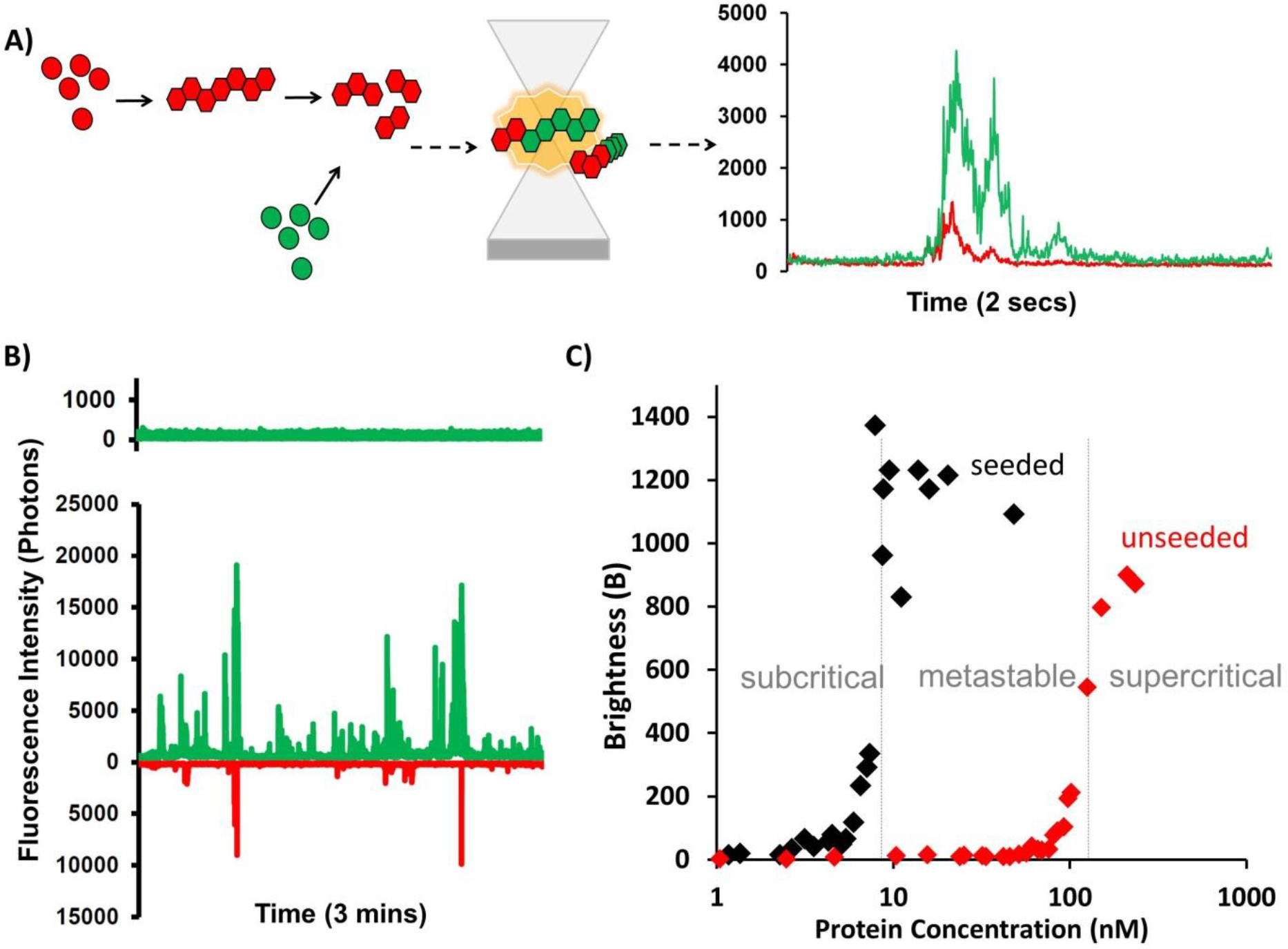
MyD88 polymerises in a concentration-dependent manner and can be self-seeded. (A) Schematic diagram of the principle of two-colour seeding experiments testing the self-replication propensity of full-length MyD88 filaments. Full-length MyD88 is expressed in mCherry-tagged version above supercritical concentration to create filaments, which are gently spun and washed, then sonicated to increase the number of fragments. These “seeds” are then mixed in a sample expressing GFP-tagged full-length MyD88 at sub-critical concentrations. (B) Example of fluorescence time trace for MyD88 at 10nM concentration. Unseeded sample demonstrating a monomeric time trace profile (above) with the seeded sample (below) showing polymerisation of GFP-MyD88 upon the addition of MyD88 ‘seeds’. (C) The B parameter (brightness) correlating with number of oligomers detected in typical time-traces as a function of protein concentration (nM), with and without “seeds” introduced. Subcritical, supercritical and “meta-stable” zones labelled.

This analysis was conducted in a systematic manner, at all of the concentrations previously described. Figure 3C shows that seeding of MyD88 polymerisation is occurring throughout a large range of concentrations. This is particularly obvious within the range of sub-threshold concentrations, where spontaneous polymerisation does not take place. Ultimately, this allowed us to define three behaviours corresponding to three zones of protein concentrations. In the subcritical zone (<10nM), full-length MyD88 does not polymerise, even in the presence of seeds. In the supercritical zone (>120nM), full-length MyD88 can spontaneously aggregate, but addition of seeds increases the effect and the plateau is reached earlier. A large metastable zone (>10-<120nM) exists where the tendency of MyD88 to aggregate on its own is low but polymerisation is catalysed by the presence of a seeding event. The presence of these three well-defined zones leads us to believe that the large protein filaments forming were not just mere inert aggregation events but a dynamic and structured process.

Biologically, the existence of this metastable zone is important, as it shows that rapid amplification of MyD88 signalling can be achieved through seeding. The *‘in vitro’* seeding is the introduction of MyD88 filaments; however, *in vivo*, seeding can be triggered by upstream proteins such as the recruitment of MyD88 through Mal nucleation. The depth of the metastable zone is also important: if this zone is too narrow, the system would respond too fast, initiating the highly effective pro-inflammatory innate immune response. A large metastable zone is therefore more physiologically desirable^24^.

### Disease-associated point mutations abrogate MyD88s ability to optimally polymerise

Having established that full-length MyD88 can undergo an active polymerisation process, we then asked whether pathological point mutations could affect this protein polymerisation propensity. Hence, L93P, R196C and L252P point mutations were separately cloned into the GFP-tagged wild-type full-length MyD88. Once again, the cell-free translation system was used and fluorescence time traces were measured and plotted as distributions of fluorescence intensities.

As shown in Figure 4, all mutations significantly reduce the ability of MyD88 to form large polymers. The example fluorescence time traces in Figure 4A as well as the FCS data clearly demonstrate the diminished polymerisation and reduction in sizes of the protein species (Figure 4B). Brightness profiles of the mutants were compared to those obtained for the isolated domains (Figure 4C,D,E). The L93P mutant polymerisation profile roughly mimics that of a MyD88 TIR domain alone, while R196C and L252P mutants polymerisation profiles approximately mirror that of a MyD88 DD alone. It appears that the point mutations decrease the capacity of the domains to contribute to polymerisation, possibly through impairing homotypic protein-protein interactions (PPIs).

**Figure 4:**
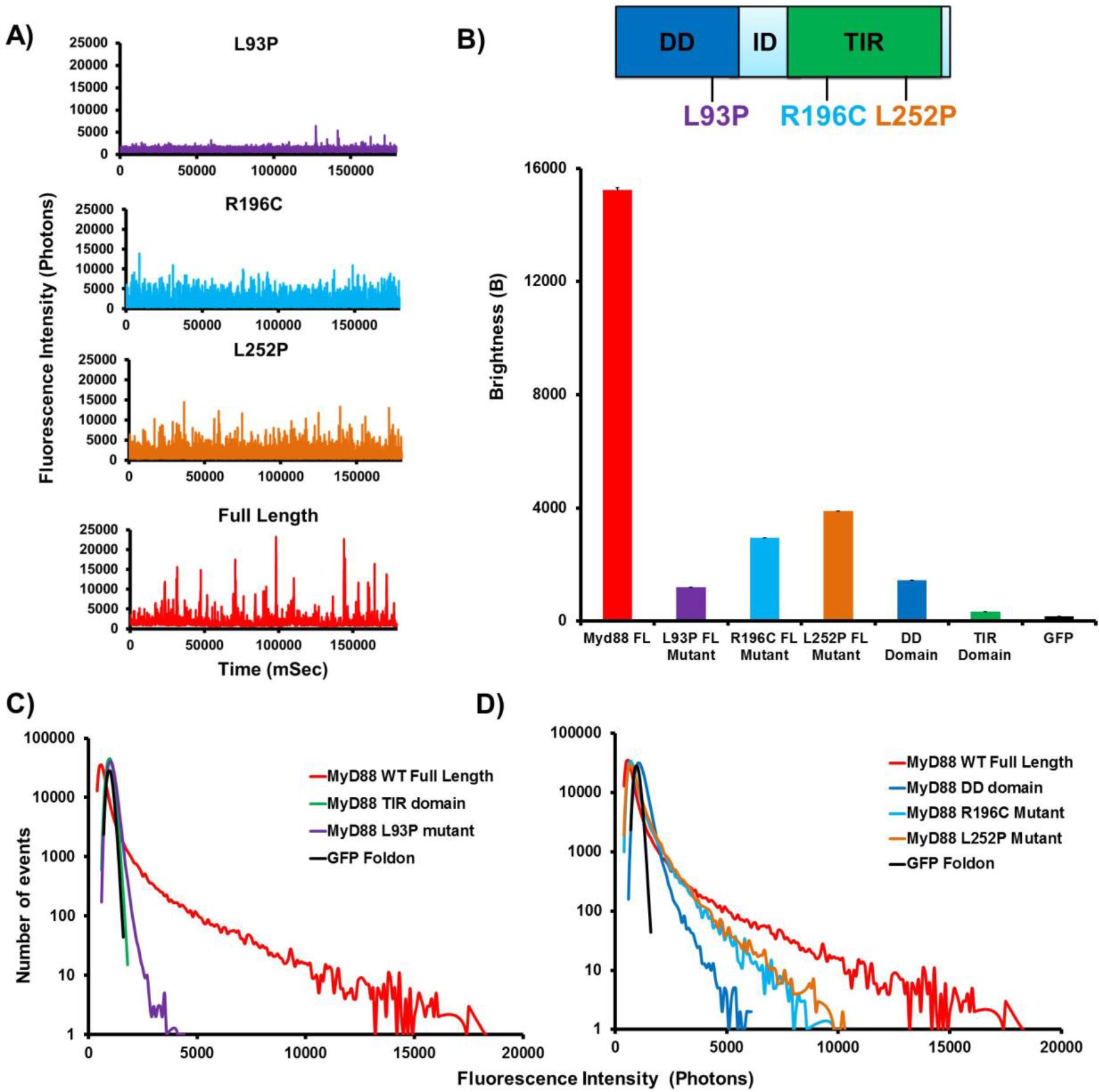
Disease-associated point mutations abrogate domain function and thus, MyD88 polymerisation. (A) Fluorescence time-traces obtained with disease-associated point mutants of the full-length MyD88 protein, as well as wild-type full-length MyD88. As in Fig. 1, the diffusion of an oligomer equates to the same number of fluorophores moving through the confocal volume, creating a burst of fluorescence in the time-trace being directly proportional to the size of the oligomer. (B) Brightness histogram showing the drastic reduction in polymer formation in comparison to the wild-type protein. (C) Fluorescence intensity histogram showing that the L93P point mutation, which is within the DD, in GFP-tagged MyD88 renders the polymerisation propensity similar to the MyD88 TIR domain alone. (D) Fluorescence intensity histogram demonstrating that the R196C and L252P point mutations (present within the TIR domain) in the GFP-tagged MyD88 render the polymerisation propensity more similar to the MyD88 DD alone.

### L252P mutants form stable oligomers at a 40-fold lower concentration than wildtype MyD88

We once again decided to examine the mutants’ behaviour as a function of protein expression, taking advantage of the control one can exert with the cell-free translation system. Figure 5A demonstrates the differences in polymerisation profiles exhibited by the point mutations at the same low concentrations (3nm). Just as in the case of wild-type MyD88, the oligomerisation thresholds of the mutant MyD88 proteins were measured and analysed by plotting the B parameter as a function of protein concentration (Figure 5B). In the case of R196C and L93P, the B values never reach those of the wild-type, indicating that the pathological point mutants cannot propagate polymerisation, no matter what concentration of protein is utilised. Mutant L252P also never forms the large aggregates that are observed with MyD88 wild-type. Strikingly though, at very low concentrations, where wild-type MyD88 and the other disease-associated point mutants exist only as monomers, L252P can still form stable low order oligomers (Figure 5B). The threshold for oligomerisation is extremely low (>2 nM). Interestingly, this threshold of assembly correlates with the concentration about which MyD88 can be seeded (Figure 3C), confirming the hypothesis that L252P acts as an activated form of MyD88^7^. The presence of these oligomers had been postulated previously based on from computational model studies^25^, hinting at their existence as oligomers at levels that are physiologically present even without expression upregulation upon receptor-ligand binding and activation. In conclusion, we have confirmed the presence of these extremely stable low-order oligomers of MyD88 and hypothesise that although wild-type full-length MyD88 is the main propagator of exponential signalling, a stable low-order oligomer formed due to mutation may be all that is needed for constitutive signalling and activation of the pathway to occur, leading to the progression of cancer.

**Figure 5:**
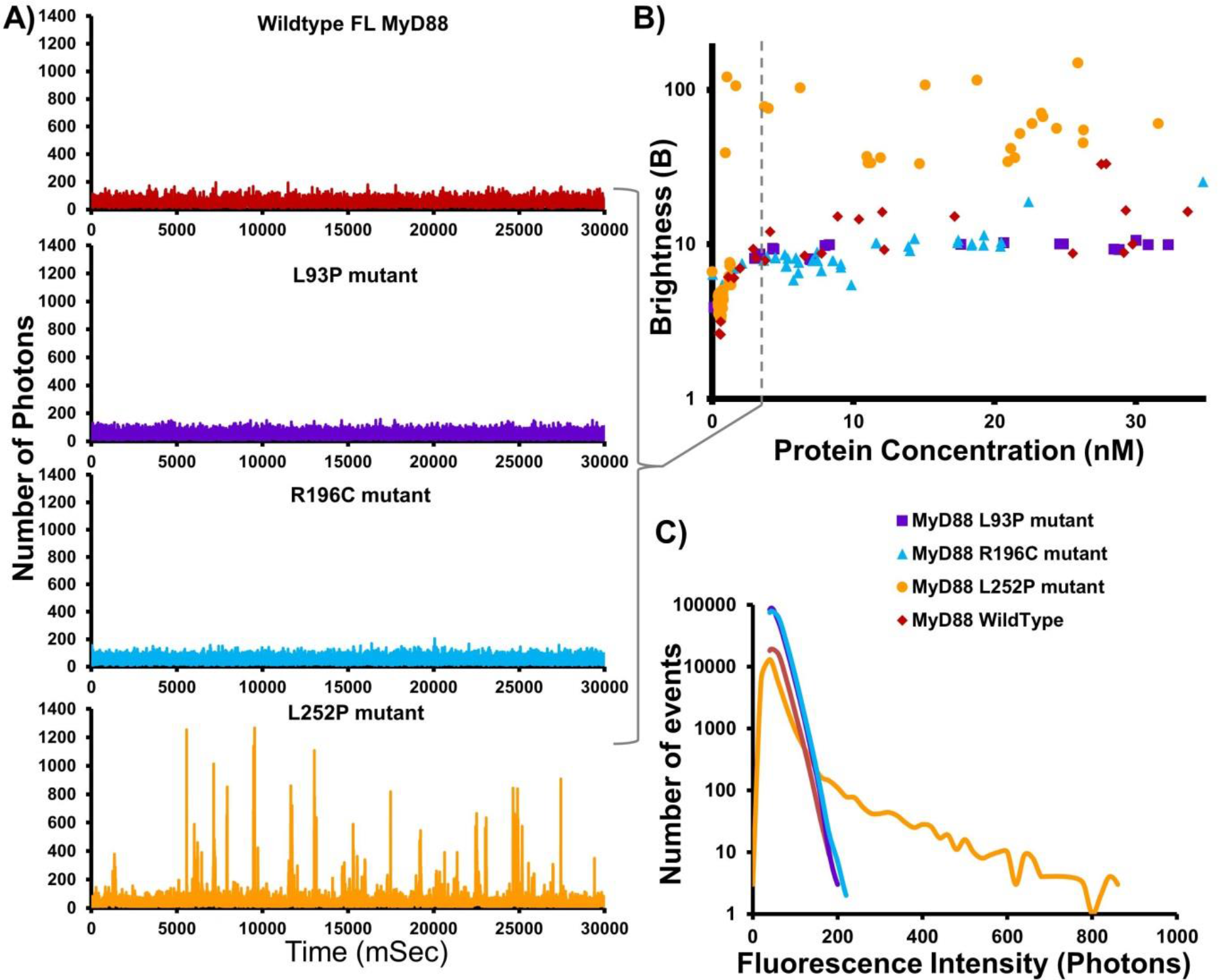
Mutations in same domain lead to contrasting disease phenotypes; cancer-causing L252P mutation lowers threshold for MyD88 oligomerisation. (A) Fluorescence time-traces obtained at ^~^3 nM protein concentration of the disease-associated point mutants in the full-length MyD88 protein, as well as full-length wild-type MyD88, demonstrating the stability of the L252P point mutation. (B) The B parameter (brightness) correlating with number of oligomers detected in typical time-traces as a function of protein concentration (nM). (C) Fluorescence intensity histogram demonstrating the stable L252P oligomer still forming at ^~^3 nM in comparison to the other constructs.

### Mutations within the same domain can lead to contrasting protein properties

Our data also show a drastic difference in the behaviour between L252P and R196C mutants, even though both residues are located within the same TIR domain. Differences in oligomerisation pattern could explain the differences in the related pathologies with L252P oligomerisation leading to cancer while R196C’s lack of polymerisation propensity dampening the innate immune response to bacterial infection. However, these are not the only differences uncovered between these disease-associated mutants. The L252P mutation is a dominant mutation, whereas L93P and R196C are both recessive mutations. Since primary immunodeficiency only affects homozygous or compound heterozygous carriers of point mutations L93P and R196C, we hypothesised that polymeric propagation could be rescued in the presence of the wild-type protein. To test this, GFP-tagged mutants and mCherry-tagged wild-type MyD88 were co-expressed in LTE and subjected to our brightness assay. The brightness parameters for the mutants obtained through single or co-expression could then be compared (Figure 6A). In the case of L93P and R196C, the brightness value is significantly larger upon co-expression, indicating that higher-order polymers were forming. Indeed, examination of the fluorescence time traces reveals the presence of coincident peaks (Figure 6B- D), showing that wild-type MyD88 can recruit the mutants into its polymers. The overall degree of polymerisation is still lower than in the case of wild-type protein alone, but the ability of the system to form large objects may be sufficiently restored to allow normal signalling to occur. In contrast, the brightness of L252P upon co-expression is unchanged in Figure 6A. Furthermore, few coincident peaks were detected (Figure 6D), indicating that the mutant species oligomerises regardless of whether the wild-type is present and that wild-type MyD88 does not seem to recruit this mutant into its polymers as readily as with L93P and R196C (Figure 6B-C). It should be noted that we have observed an intriguinging codiffusion pattern for the R196C MyD88 mutant, with the coincidence peaks exhibiting inverse relationship patterns suggesting that the filament is made up of distinct stretches of mutant protein and wildtype MyD88 species cofibrillating within the fibril (Figure 6C). The differential incorporation of the mutants into the wild-type polymers correlates well with what is observed at the physiological level. Heterozygous patients carrying the L93P or R196C mutations do not suffer from the recurrent bacterial infections. The wild-type protein polymerisation, as well as the incorporation of the mutants into the polymerising wild-type protein, albeit at suboptimal levels (Figure 6A), may be sufficient to propagate signalling efficiently. Impairment of polymerisation and subsequent signalling is only observed in the absence of wild-type MyD88, as would be the case for homozygous and compound heterozygous carriers (i.e. both alleles of the gene harbour mutations such as L93P and R196C)^4, 5^. L252P is not observed to incorporate with the wild-type MyD88 polymer, with our data suggesting the maintenance of a distinct population of finite sized oligomers, irrespective of the presence of wild-type (Figure 6A&D). This would correlate with the fact that both the heterozygous and homozygous patients suffer from cancer^26^.

**Figure 6:**
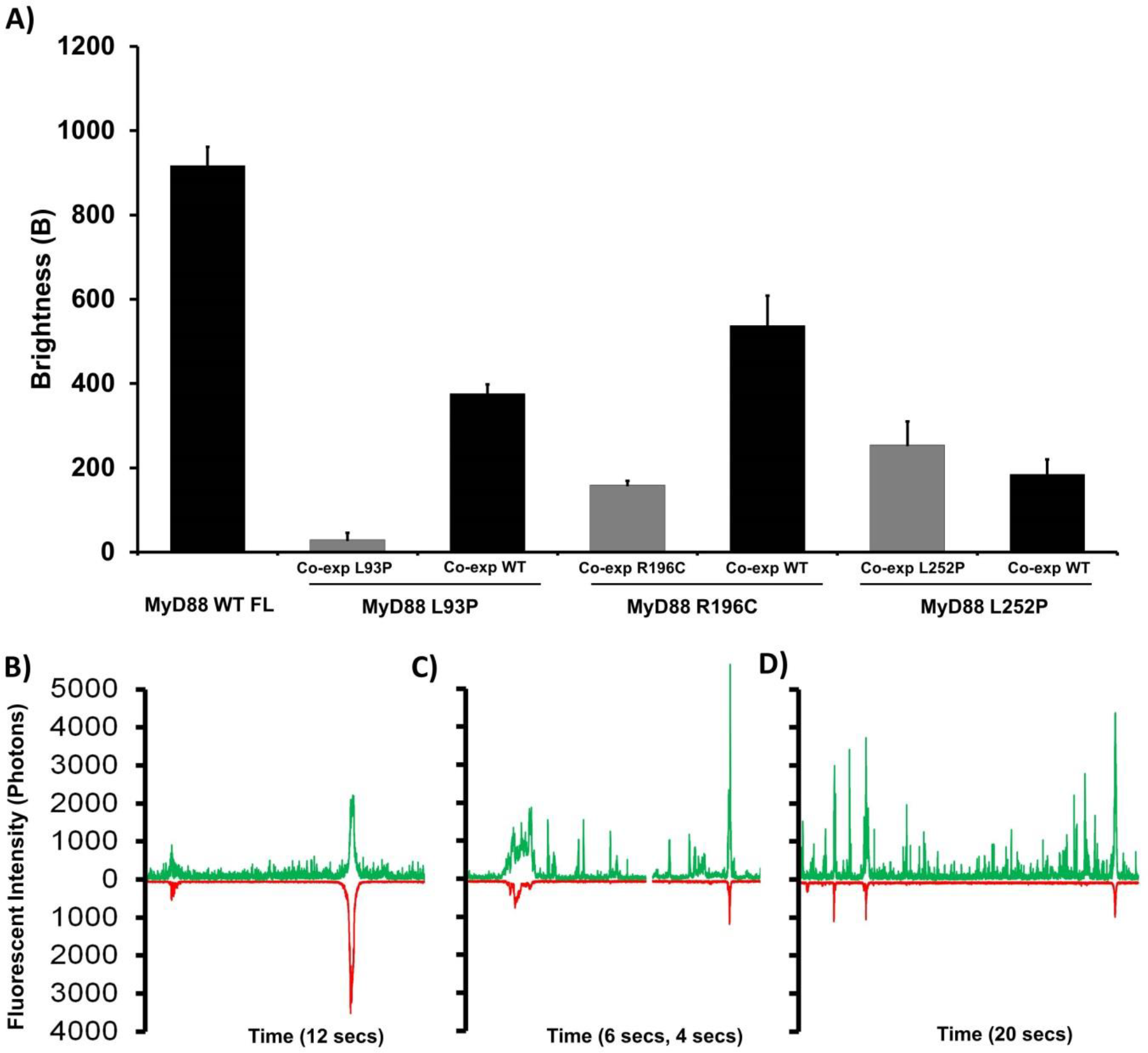
Co-expression with wild-type full-length MyD88 partially rescues the ability of the recurrent bacterial infection disease-associated point mutants to polymerise. (A) Brightness histogram of the mCherry-tagged wild-type MyD88 co-expressed with disease-associated mutants (simulating heterozygous expression in patients), as well as L93P, R196C or L252P mutant proteins co-expressed with themselves (i.e. homozygous protein expression) and wild-type MyD88 alone as a control. (B-D) Fluorescence time traces of the disease-associated mutants co-espressed with mCherry-tagged wild-type MyD88. The recurrent bacterial infection disease-associated point mutation, L93P (B) and R196C (C), co-expression rescue experiments contrast with the continuously oligomerising L252P mutant (D) whereby L252P is not incorporated / rescued and exists as its own separate population.

## DISCUSSION

### Full-length MyD88 biophysical behaviour and its link to signalling

Here, we studied the contribution of the domains and the effect of physiological mutations on the biochemical and biophysical behaviour, in particular on the polymerisation propensity of MyD88, a key protein in the TLR pathways. To characterise the formation of protein assemblies, we utilise single molecular fluorescence spectroscopy, as this technique has the unique ability to quantify oligomers and track conformational changes at a single protein level. Through utilising cell-free eukaryotic expression, we can co-express proteins together in their known complexes, allowing the native and physiological PPIs to occur. We can also control expression and therefore, distinguish thresholds, aggregation propensity, and self-propagating behaviour.

When we first compared the isolated domains with the full-length protein, we demonstrated that only full-length MyD88 is capable of forming large objects in a concentration-dependent, self-templated manner. Traditionally, the biochemical studies of MyD88 and other adaptors have mainly focussed on the role of the isolated domains, partly due to the difficulty to purify the full-length proteins. From the study of these isolated domains, two mechanisms of self-assembly have been described. For many years, TIR domain associations, which are weak and transient, have been seen to contribute little to the oligomerisation status of signalling proteins^27, 28^. However, the recent cryoEM structure of Mal TIR domain in filamentous form, as well as novel studies^12^,demonstrated that upon specific ligand binding TIR-containing proteins cooperatively assemble into large multi-protein complexes^12, 23^. The TIR domain of MyD88 was also demonstrated to polymerise but only upon seeding by Mal filaments. On the other hand, the DD has been known for participating in the formation of the high-order helical assembly of the MyDDosome, a signalling complex that also includes the DD of IRAK2 and IRAK4. Our data using the isolated domains recapitulate those findings (Figure 2). We show that the TIR domain alone are present as small, low-order oligomers, no matter what concentration of protein is expressed and that these TIR domain oligomers never combine to form large signalosomes on their own (Figure 1 & Figure 2). In our system, MyD88 DD is able to form oligomers of approximate octomeric sizes, consistent with previous results^11^. The DD can form larger assemblies though these events are very rare (data not shown). Interestingly, the DD appears to exhibit concentration-dependent ‘all-or-none’ monomeric to oligomeric behaviour, albeit at a much lower scale that the full-length MyD88 (Figure 1B inset). The behaviour of the isolated domains is in stark contrast to the full-length protein, which has a tendency to form large complexes at these low concentrations (Figure 1). Both the TIR and death domains can drive oligomerisation; they are necessary but not sufficient for polymerisation. Our seeding assay also reveals a large zone of concentrations where the full-length protein is metastable. We hypothesize that the presence of the two domains contributes to the creation of this metastable zone. Having two domains has been demonstrated to create auto-inhibition within full-length proteins so as to hinder spontaneous assembly. This can be extrapolated from our data when comparing non-seeded full-length MyD88 with TIR only or DD only threshold concentrations (Figure 2), therefore complementing the model that having two domains stabilizes polymer formation for a delayed, yet faster response^24^.

This new biophysical data incorporates well into the existing model of signalling. Studies agree upon the necessity of Mal oligomerisation for the propagation of the signal upon receptor activation. Yet, in cells, full-length MyD88 or its DD overexpression can overcome the need for Mal oligomerisation. The prion-like polymerization of MyD88 would explain this feature. MyD88 full-length or DD alone have the potency to oligomerise in a higher-order manner on their own but at higher concentration (as in the case of overexpression). However, polymerisation has the potential to occur at endogenous levels of MyD88 in the presence of a ‘seed’. This suggests that Mal oligomerisation acts as a nucleation platform for MyD88 as has been put forward in the present published models^29^. MyD88 then recruits downstream death domain (DD)-containing proteins through homotypic interactions to form a single left-handed helical scaffold, held together by 3 conserved DD interaction types^30^. MyD88 polymers serve in turn as scaffolds for the IRAK kinases, allowing protein binding, signal propagation, amplification and ultimately, rapid signal transduction.

### Pathological point mutants reveal the importance of oligomerisation in signalling

We also characterised the propensity of three pathological mutants to self-interact and to create. Our results correlate with the various disease phenotypes observed, as each mutant perturbs the PPIs networks in a different manner.

Previous studies have uncovered the impact of disease-associated point mutations of MyD88 in regard to the heterotypic protein-protein interactions that occur with the other components of the signalling pathway^29, 31,4^. The MyD88 assembly is composed of two parallel strands of MyD88 TIR subunits; (Figure 7, strand I (cyan) and strand II (orange)). The interactions between MyD88 subunits within each strand (intrastrand interactions) involves opposite sides of the TIR domain with residues in the BB loop (“BB surface”) of one subunit interacting with residues on the “EE surface” on the next subunit. R196 is part of the BB loop and is predicted to stabilize the conformation required for assembly formation by forming a salt bridge with E183 in the αA helix. The R196C mutation is likely to destablise the BB loop conformation. The R196C mutant was found to have reduced PPIs with other TIR domain-containing signaling proteins^4^. Recently, Ve et al. demonstrated that an R196 mutant completely abolished Mal TIR-induced assemblies of MyD88 TIR, as well as the ability of full-length MyD88 to cluster in HEK293 cells, confirming the crucial role of this amino acid in TIR homotypic binding. We wished to uncover the impact of the R196C mutation in regard to the homotypic interactions that underly the self-association of full-length MyD88. Consistent with previous work, our data demonstrates that the R196C mutant has a reduced ability to homotypically interact and therefore polymerise compared to the wild-type protein.

**Figure 7:**
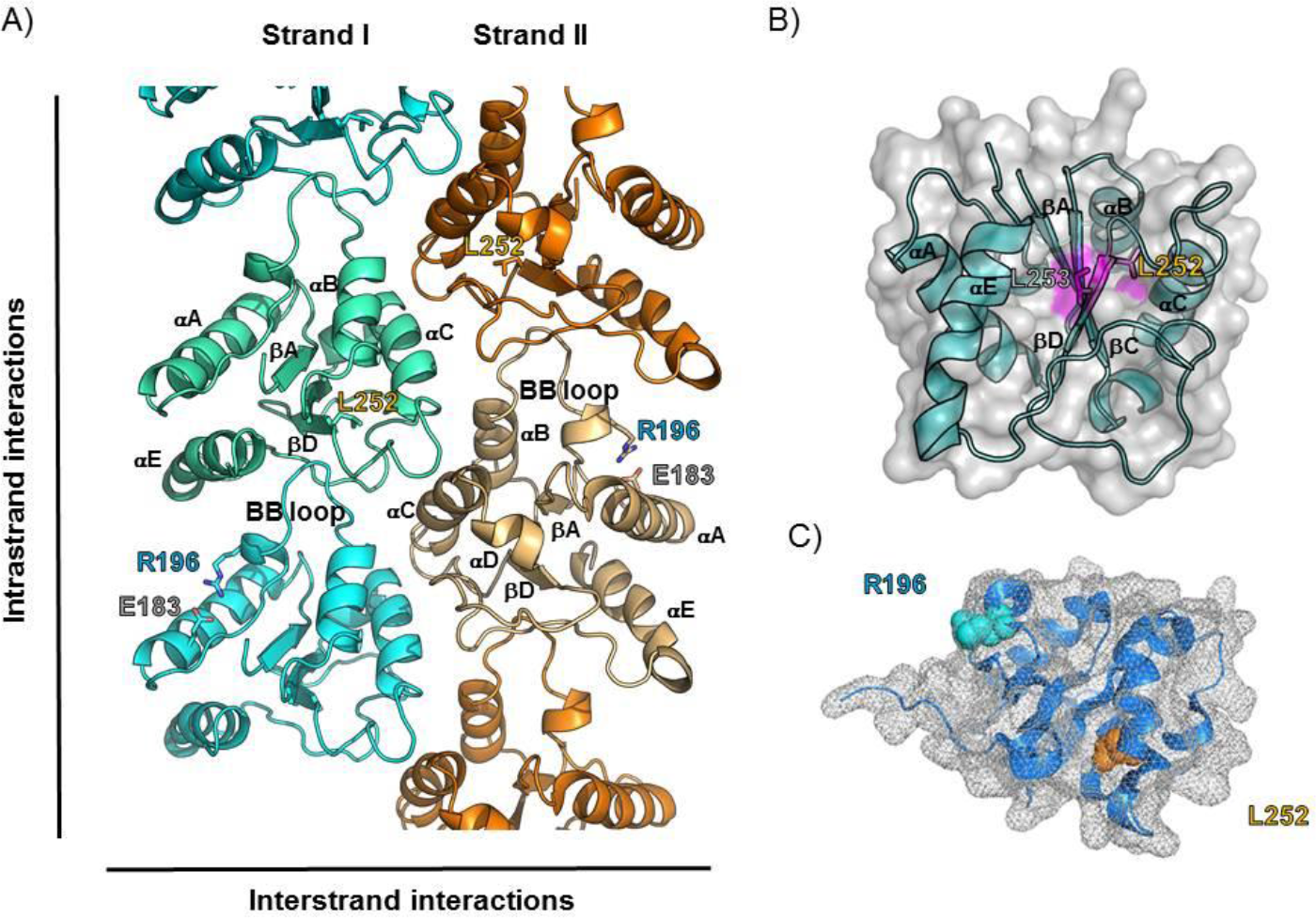
MyD88 TIR domain assembly model and mapping of disease-associated mutations. (A) Cartoon representation of a MyD88TIR domain assembly model based on the MALTIR filament structure with mutants highlighted^12^.(B) Surface/cartoon representation of the “EE surface” (the αE helix, βD strand, βE strands) of one MyD88TIR subunit, L252 mutant highlighted (orange). (C) Structure of the MyD88 TIR domain (PDF ref:4eo7), in which the R196 was pinpointed in cyan and the L252 in orange.

The L93P mutation in the death domain (DD) has been shown to prevent interactions with the downstream proteins of the signalling cascade such as IRAK4^4^. As the highly conserved L93 side chain is buried, the L93P mutation, as well as affecting the helix formation, disrupts the hydrophobic core of the DD^9^. The view is that this point mutation renders the DD non-functional and prevents the optimal binding to upstream signalling proteins, as well as fully abrogating binding to downstream signalling proteins, such as the kinases like IRAK4 that propagate the signal. Again, we aimed to uncover the effects of this mutation upon the homotypic interactions of the full length MyD88 protein. We discovered a reduced propensity for homotypic interactions between MyD88 itself, with L93P unable to form polymers to the same extent as wild-type MyD88. Although L93P and R196C occur in the two different domains of MyD88, one in the DD and one in the TIR domain respectively, they both cause autosomal recessive MyD88 deficiency that results in life-threatening, recurrent pyogenic bacterial infections. We have established two molecular links between the mutants by showing that they both exhibit a reduced ability to polymerise and that they can both be partially incorporated (“rescued”) by the presence of the wild-type protein. This may explain the recessive character of the disease, as only homozygous or compound heterozygous carriers display the disease phenotype.

On the other hand, it intrigued us that although both R196C and L252P occur in the same TIR domain, one gives rise to recurrent bacterial infections associated with the immunodeficiency, while the other results in lymphoma. L252 is the first residue of the BD strand and is located next to the highly conserved I253 residue of the EE surface, as can be seen in Figure 7. The L252 sidechain is largely buried and does not contribute directly to the interaction with the BB loop of the preceding MyD88 subunit in the assembly, but a mutation of this residue to a proline (L252P) is likely to induce conformational changes resulting in a “EE surface” with different properties. So far, computational methods have been mainly used to characterize the conformational effects of the L252P mutation^7, 32, 33^. Molecular dynamics simulations revealed that the L252P mutation allosterically quenched the global conformational dynamics of the TIR domain and readjusted its salt bridges and dynamic community network. The dampened motion restricts its ability to heterodimerize with other TIR domains, thereby curtailing physiological signalling. Interestingly, results indicate that the mutation stabilizes the core of the homodimer interface of the MyD88-TIR domain^25^. This would increase the population of homodimer-compatible conformational states in MyD88 family proteins, enhancing signalling. Our experimental results confirm, for the first time, that L252P forms extremely stable oligomers compared to the wild-type protein and the other mutants we have studied. We could observe oligomers when the protein was expressed at concentrations as low as approximately 2nM. As suggested, the conformational dynamics of cancer-associated MyD88-TIR domain mutant L252P allosterically seem to tilt the landscape toward homo-oligomerization *in vitro*, which would quench physiological signaling *in vivo*, propagating a signal independently of the TLR ligands^33^.

## CONCLUSION

Our observations suggest that prion-like polymerisation is a fundamental mechanism of intracellular communication and drives the complex interplay that exists among the major signalling cascades of the innate immune system. Our disease-associated mutant protein data highlight the crucial functions that these large signalling platforms have in the optimal execution of defence against pathogens and intracellular homeostasis, regulating the critical first line of defence that is the innate immune system. Development of drugs that can interfere with the higher-order oligomerisation and polymerisation of adaptor proteins would therefore be a novel advancement in medicine, with the potential to function as anti-inflammatory as well as anticancer agents. The role of MyD88 in cancer is now emerging. MyD88 L252P/ L265P is implicated in almost 100% of Waldenström’s macroglobulinemia (WM) cases, 2–10% of chronic lymphocytic leukemia (CLL) cases, 69% of cutaneous diffuse large B cell lymphoma (DLBCL) cases, and 38% of primary central nervous system lymphomas (PCNSL) cases ^32^. The activated B-cell-like (ABC) subtype of diffuse large B-cell lymphoma (DLBCL) remains the least curable form of this malignancy, with less than a 40% cure rate^34^. Despite the expanding knowledge on MyD88 since 2012, the L252P mutation of MyD88 was identified in tumour samples from 49 of 54 patients with the incurable form of the disease. Finally, since 2016, testing for MyD88 was added to the essential recommendation2019;s Macroglobulinemia (LPL/WM) in the National Comprehensive Cancer Network (NCCN) Guidelines. The strong link between the mutation and cancer makes the stable signalling oligomers created by MyD88 L252P an enticing target from a therapeutic standpoint.

## MATERIALS and METHODS

### Preparation of LTE

Cell-free lysate was collected from *Leishmania tarentolae (LT)* as per Johnston & Alexandrov ^17, 35, 36^. *Leishmania tarentolae* Parrot strain was acquired as LEXSY host P10 from Jena Bioscience GmbH, Jena, Germany and cultured in TBGG medium containing 0.2% v/v penicillin/streptomycin (Life Technologies) and 0.05% w/v hemin (MP Biomedical). LT cells were harvested through centrifugation at 2500 × *g*, washed twice by resuspension in 45 mM HEPES (pH 7.6) containing 3 mM magnesium acetate, 100 mM potassium acetate and 250 mM sucrose. Cells were resuspended to 0.25 g cells/g suspension and incubated under 7000 kPa nitrogen for 45 minutes, then lysed by rapid release of pressure in a cell disruption vessel (Parr Instruments, USA). Through sequential centrifugation at 10,000 ×*g* and 30,000 ×*g*, the cell-free lysate was clarified and 10 μM anti-splice leader DNA leader oligonucleotide was added. The cell-free lysate was then desalted into 45 mM HEPES (pH 7.6) containing 100 mM potassium acetate and 3 mM magnesium acetate. The LTE was supplemented with a coupled translation/transcription feeding solution and snap-frozen until required for further experimentation.

### Gateway plasmids for cell-free protein expression

Full-length MyD88, MyD88 TIR domain(159-296a.a), MyD88 DD (1-117a.a) as well as all of the other proteins from the pathway were cloned into the Gateway destination vectors: N-terminal GFP-tagged (pCellFree_G03), N-terminal mCherry-tagged (pCellFree_G05), C-terminal eGFP-tagged (pCellFree_G04) or C-terminal mCherry-cMyc-tagged (pCellFree_G08), facilitating cell-free expression^37^. The Gateway PCR cloning protocol was used and entry clones were generated with PCR primers to attB1 and attB2 sites (forward primer: 5’GGGGACAAGTTTGTACAAAAAAGCAGGCTT (nnn)_18-25_ 3’, reverse primer: 5’GGGGACCACTTTGTACAAGAAAGCTGGGTT (nnnn)_18-25_ 3’)^38^.

### Cloning point mutations

Primers were designed and ordered through IDT. Cloning was conducted as per Phusion^®^ High-Fidelity DNA Polymerase protocol with full-length MyD88 N-terminal GFP tagged (pCellFree_G03) and C-terminal mCherry-cMyc tagged (pCellFree_G08) as donor construct. All mutants sequence were verified by Ramiciotti UNSW Sequencing facility.

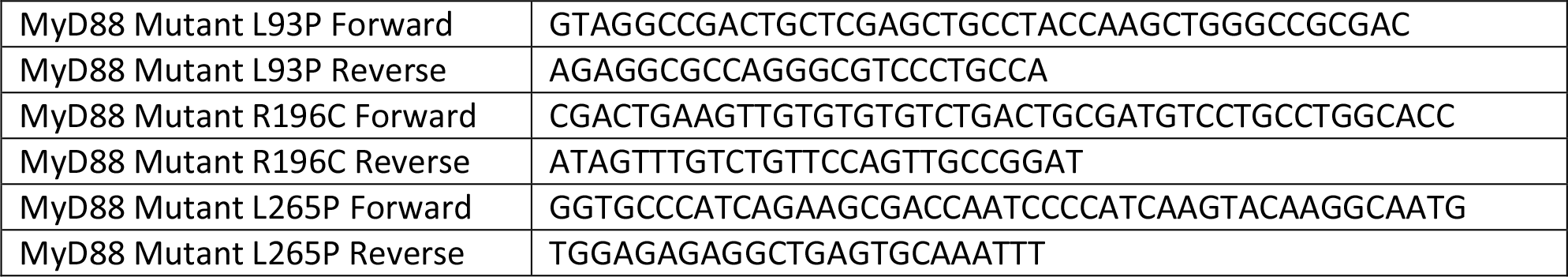

### *In vitro* protein expression

All proteins were expressed through the addition of LTE lysate to DNA template (in a ratio of 1 :9) for h at 27 °C and 0.5 h at 37 °C. For protein dilution titrations, the concentration of DNA template was varied using serial dilutions with nuclease-free H_2_0 covering ranges from 600 nM stock to 50 nM concentrations correlating with protein concentrations of 30 nM to 0 nM. 1 μL of the diluted DNA was used to prime 9 μL of LTE. Samples were processed immediately for either fluorescence microscopy analysis, seeding experiments or AlphaScreen assay.

### Single-molecule fluorescence spectroscopy

Single-molecule spectroscopy was performed as described in previous studies by Sierecki et al. and Gambin et al.^18, 21^. The proteins were labelled with genetically encoded fluorophores (GFP and mCherry) facilitating fluorescence spectroscopy under a confocal microscope directly in the cell-free expression mixtures, without any purification steps. Two overlapping lasers excite the GFP and mCherry fluorophores, creating a small detection volume in which GFP and mCherry fluorescence emitted by proteins is recorded on single-photon-counting detectors. Due to Brownian motion the proteins freely diffuse, constantly entering and exiting the detection volume of the microscope and creating fluctuations in the fluorescence intensity. The number of photons collected versus the time of the measurements is obtained as raw data. Then the amplitude and frequency of the fluorescence fluctuations are quantified to characterize the oligomerisation status of the proteins^39^.

N-terminal trimeric foldon and GFP-foldon (both known to be trimeric proteins and used as a known size control) were expressed for quantification of the intensity measurements. A 488 nm laser beam was focused in the sample volume using a 40× / 1.2 NA water immersion objective (Zeiss). The fluorescence of eGFP was measured through a 525/20 nm band pass filter, and the number of photons collected in 1 ms time bins (*I*(*t*)) was recorded. The proteins were diluted 10 times in buffer A.

The fluorescent time-trace I(t) obtained shows the presence of intense bursts of fluorescence, with values well over the typical fluctuations of I(t). The presence of these bursts increases the standard deviation of the distribution. To compare the aggregation at different concentrations, we used the B parameter, this being independent of the protein concentration and can be written as:

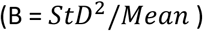

### Single-molecule fluorescence spectroscopy: seeding experiments

In this assay, full-length MyD88 was expressed as an mCherry-tagged protein. The aggregates were spun down and sonicated, and then added to a solution of monomeric GFP protein. We directly detected the recruitment of the GFP monomer to the seed with two-colour coincidence measurement, by detecting the simultaneous presence of a signal in the GFP (green) and mCherry (red) channels. mCherry-tagged seeds of full-length MyD88 were expressed in LTE by the addition of the template DNA in 10 μL lysate, generating a final concentration of ^~^30 nM protein. To produce the seeds, the samples were then spun down at 13,000 × *g* for 5 min. 80% of the supernatant was discarded, and the solution was sonicated for 1 min in a water bath. During sonication, GFP-tagged full-length MyD88 was expressed, as previously described using serial dilutions to generate a range of GFP-tagged protein concentrations from ^~^25 nM to 0 nM and diluted ten times before being placed under the microscope. Two lasers (488 nm and 561 nm) were focused in solution with a 40×/1.2-NA water-immersion objective (Zeiss). Fluorescence was collected and separated with a 565-nm dichroic mirror; signal from GFP was passed through a 525/20-nm band-pass filter, and fluorescence from mCherry was filtered by a 580-nm long-pass filter. The fluorescence of the two channels was recorded simultaneously in 1-ms time bins. Fluorescence time traces were recorded for 180 s.

### Single-molecule fluorescence spectroscopy: co-expression experiments

MyD88 point mutations with N-terminal tagged eGFP were co-expressed with full-length MyD88 C-terminal tagged mCherry-cMyc in the respective ratios of 20 and 40 nM of DNA template, in 10 μL of LTE for 2.5 h at 27 °C and 0.5 h at 37 °C, then measured on the microscope and analysed as per all single-molecule spectroscopy experiments described above.

All proteins were run on NuPAGE™ 4-12% Bis-Tris Protein Gels, 1.0 mm, 10-17 well gels from Invitrogen and imaged by BioRad.

## ABBREVIATIONS

MyD88: : Myeloid Differentiation primary response 88
TIR: : Toll/Interleukin-1 Receptor
DD: : Death Domain
ID: : Intermediate Domain
PYD: : Pyrin Domain
CARD: : Caspase Recruitment Domain
TLR: : Toll-Like Receptor
PRR: : Pattern Recognition Receptor
ASC: : Apoptosis-associated speck-like protein containing a CARD
MAVS: : Mitochondrial Anti-Viral Signalling Protein
IRAK: : Interleukin-1 Receptor Associated Kinase
GFP: : Green Fluorescent Protein
LTE: : *Leishmania tarentolae* extracts
WT: : Wild-Type
FL: : Full-Length
PPI: : Protein-Protein Interactions
FCS: : Fluorescence Correlation Spectroscopy
PCH: : Photon Counting Histogram
B: : Brightness

## AUTHOR’S CONTRIBUTIONS

AOC and BC carried out the cell-free experiments. BC cloned mutant proteins. DH and AB contributed the cell-free reagent and other reagents. AOC, BC, JB, AM and JC performed single-molecule experiments, Brightness, PCH, FCS and coincidence analysis of data. TV and BK contributed Figure 7. ES, YG and AOC designed the study. AOC, YG, ES drafted the manuscript; all authors read and approved the final manuscript.

